## Supplementary File for "Model predictive game control for personalized and targeted interactive assistance"

### Supplementary File
### Model predictive game control for personalized and targeted interactive robotic assistance

Abdelwaheb Hafs, Anaïs Farr, Dorian Verdel, Olivier Bruneau,
Etienne Burdet and Bastien Berret

In this supplementary note, we present a general dyadic affine-quadratic case that can be used to model more complex interaction dynamics. Specifically, when the actual states and inputs of the robot and human systems are modeled along with their physical connection (e.g., as spring-damper elements), the inputs from the robot and the human will influence different states of the coupled human-robot system.

As in the main text, the control $\mathbf{u}(t)$ combines the inputs from the human agent $h$ and robot agent $r$. More explicitly, we assume that $\mathbf{u}(t)$ can be written as:

$$\mathbf{u}(t) = \mathbf{S}_r \mathbf{u}_r(t) + \mathbf{S}_h \mathbf{u}_h(t)\,, \tag{1}$$

where $\mathbf{S}_r \in \mathbb{R}^{m \times m_r}$ and $\mathbf{S}_h \in \mathbb{R}^{m \times m_h}$ are reshaping matrices, with $\mathbf{u}_r(t) \in \mathbb{R}^{m_r}$ and $\mathbf{u}_h(t) \in \mathbb{R}^{m_h}$. In general, the dimensions of the robot and human inputs may differ.

In the simplest case where $m = m_h = m_r$ with perfect kinematic alignment between the robot and the human, one would define $\mathbf{S}_r = \mathbf{S}_h = \mathbf{I}$ (identity matrix) and $\mathbf{u} = \mathbf{u}_r + \mathbf{u}_h$, as in the main text. In this case, the two rigid body dynamics are considered together, and the interaction torque is canceled. Similar matrices can be defined when focusing solely on the robot's perspective and attributing the interaction torque to the human input, as seen e.g. in [1]. This approach may be relevant when carrying a large load, such that the human dynamics can be neglected. However, the reshaping matrices $\mathbf{S}_r$ and $\mathbf{S}_h$ can be used for modeling more complex physical interaction considering joint misalignments and the intrinsic human dynamics.

Considering the matrix $\mathbf{B}(t) = \dfrac{\partial \mathbf{f}}{\partial \mathbf{u}}(\mathbf{x}_d(t), \mathbf{u}_d(t), t)$ from the linearization of the interaction dynamics $\dot{\mathbf{x}}(t) = \mathbf{f}(\mathbf{x}(t), \mathbf{u}(t), t)$ around a desired trajectory/control pair $[\mathbf{x}_d(t), \mathbf{u}_d(t)]$, and defining $\mathbf{B}_r(t) \triangleq \mathbf{B}(t)\mathbf{S}_r$ and $\mathbf{B}_h(t) \triangleq \mathbf{B}(t)\mathbf{S}_h$, we obtain the following local *affine human-robot interaction model*:

$$\dot{\boldsymbol{\xi}}(t) = \mathbf{A}(t)\boldsymbol{\xi}(t) + \mathbf{B}_h(t)\mathbf{u}_h(t) + \mathbf{B}_r(t)\mathbf{u}_r(t) + \mathbf{c}(t)\,. \tag{2}$$

In these settings, the Nash equilibrium can be obtained by solving the coupled Riccati-like differential equations (see Corollary 6.5 in [2], page 323):

$$
\begin{aligned}
-\dot{\mathbf{P}}_i \; &= \; \mathbf{F}^\top \mathbf{P}_i + \mathbf{P}_i \mathbf{F} + \mathbf{Q}_i + \sum_{j\in\{r,h\}} \mathbf{P}_j \mathbf{B}_j \mathbf{R}_j^{-1} \mathbf{R}_{ij} \mathbf{R}_j^{-1} \mathbf{B}_j^\top \mathbf{P}_j \,, \\
-\dot{\boldsymbol{\alpha}}_i \; &= \; \mathbf{F}^\top \boldsymbol{\alpha}_i + \mathbf{P}_i \boldsymbol{\beta} + \sum_{j\in\{r,h\}} \mathbf{P}_j \mathbf{B}_j \mathbf{R}_j^{-1} \mathbf{R}_{ij} \mathbf{R}_j^{-1} \mathbf{B}_j^\top \boldsymbol{\alpha}_j \,, \\
\mathbf{F} \triangleq \mathbf{A} &- \sum_{i\in\{r,h\}} \mathbf{B}_i \mathbf{R}_i^{-1} \mathbf{B}_i^\top \mathbf{P}_i \,, \quad \boldsymbol{\beta} \triangleq \mathbf{c} - \sum_{i\in\{r,h\}} \mathbf{B}_i \mathbf{R}_i^{-1} \mathbf{B}_i^\top \boldsymbol{\alpha}_i \,.
\end{aligned} \tag{3}
$$

This complete solution includes $\mathbf{R}_{hr}$, which could be used to model that the human seeks to minimize the robot effort (whereas in the main text we only assumed that the robot assists the human using $\mathbf{R}_{rh}$). In the main text, we instead set $\mathbf{R}_{hr} = \mathbf{0}$ and used the notation $\mathbf{R}_{hh} = \mathbf{R}_h$ and $\mathbf{R}_{rr} = \mathbf{R}_r$. In this general affine-quadratic case, the Nash-optimal control laws are given by:

$$
\mathbf{u}_i(t, \boldsymbol{\xi}(t)) = -\mathbf{R}_i^{-1} \mathbf{B}_i(t)^\top [\boldsymbol{\alpha}_i(t) + \mathbf{P}_i(t)\boldsymbol{\xi}(t)] \,, \quad i \in \{r, h\} \,, \quad t \in [\tau, \tau + \triangle_p] \,. \tag{4}
$$

The MPG control method presented in the main text can readily be applied with these modified equations.
